# Censoring-aware Gaussian process modeling identifies time informative functional dynamics associated with spoilage status of modified atmosphere packaged broiler meat

**DOI:** 10.64898/2026.06.11.731636

**Authors:** Elio Nushi, Julia Manninen, Per Johansson, Antti Honkela, Johanna Björkroth

## Abstract

Microbial spoilage of packaged meat is driven by complex microbial succession and related metabolic activity. The current tools to assess product quality and determine shelf life do not fully serve food manufacturers. Functional time informative approaches that capture the active metabolic state of the spoilage microbiome and predict the rate of spoilage are lacking. We developed a censoring-aware Gaussian process (CAGP) framework to model longitudinal pathway expression profiles of broiler meat spoilage microbiomes. We used two series of metatranscriptomic data generated in our previous study of samples stored either at 4 or 6°C. The samples had also been annotated with odor-based sensory scores defining fresh, early-spoilage, and late-spoilage phases. The 4°C time series were used to develop and evaluate the temporal prediction framework, while the independent 6°C series were used to assess whether the functional spoilage trajectories learned at 4°C could generalize across temperature conditions. The CAGP models smooth temporal trajectories of pathway activity while accounting for uncertainty and for observed zero values that may arise from non-detection rather than true biological absence. In this framework, zeros are treated as left-censored observations below a soft detection threshold. Within the 4°C time series, leave-one-replicate-out (LORO) and leave-one-day-out (LODO) analyzes predicted the sampling time of held-out samples with mean absolute errors of 0.43 and 0.78 days, respectively. In the LORO analysis, predicted spoilage phases also largely agreed with the sensory classification. When trajectories learned at 4°C were applied to the independent 6°C samples, the predicted samples retained their temporal ordering along the spoilage trajectory, suggesting the preservation of shared functional spoilage programs despite temperature-dependent differences in spoilage rates. Cross-entropy ranking further identified pathway modules carrying time- and phase-informative signals across temperatures. Overall, this framework provides a probabilistic approach for linking metatranscriptomic functional dynamics to sensory spoilage progression, supporting shelf-life assessment beyond retrospective microbial enumeration.

**Highlights:**

- Functional pathway activity tracks spoilage in refrigerated broiler meat.
- A censoring-aware Gaussian process handles non-detected pathway activity values.
- Longitudinal models predict storage time from microbial pathway activity.
- Shared pathway signals predict spoilage at two refrigerated temperatures.
- Cross-entropy scoring identified pathways linked to spoilage progression.

## 1. INTRODUCTION

Microbial spoilage of modified atmosphere packaged (MAP) meat is driven by the succession and metabolic activities of a subset of microorganisms that generate unacceptable odors and flavors over time (Borch et al., 1996; Pothakos et al., 2015; Snyder et al., 2024). Although the organisms typically associated with meat spoilage are generally known, the actively transcribed genes and metabolic pathways underlying spoilage progression remain less well understood. In particular, increasing our understanding of meat spoilage requires linking sensory deterioration with the active functional state of the microbiome as part of a dynamic process. A better understanding of the longitudinal behavior of spoilage microbiomes during shelf life and beyond may provide new possibilities for monitoring and predicting shelf life using functional molecular data.

Metatranscriptomic profiling provides a way to capture the expression of microbial metabolic pathways during spoilage and to identify functional programs associated with sensory deterioration. In our previous study by Manninen et al., 2026, spoilage phases were defined based on sensory evaluation and linked to metatranscriptomic and metabolomic data. The study showed that the microorganisms commonly considered responsible for spoilage were not necessarily all transcriptionally active, and some active taxa did not by themselves explain spoilage progression. The previous study connected pathway activity and metabolomic outcome to sensory-defined spoilage phases whereas it did not explicitly model the time-resolved dynamics of pathway expression.

Our present study extends this previous work by investigating whether longitudinal pathway-expression patterns can be used to describe and predict the progression of broiler meat spoilage over time. Rather than comparing pathway activity only between predefined spoilage phases, we model pathway trajectories across storage days and examine how these trajectories relate to the transition from fresh product to early and late spoilage. This allowed us to ask not only whether particular pathways are associated with spoilage phases, but also whether pathway dynamics contain sufficient temporal information to infer the storage time and spoilage state of individual samples. In addition, because spoilage progression is temperature-dependent, we investigated whether models trained on pathway dynamics can capture differences between samples stored at 4°C and 6°C. In the previous study by Manninen et al., 2026, observations suggested that pathway activity profiles at these temperatures are broadly similar, although metabolism and spoilage progression appeared faster at 6°C. This motivates testing whether temperature affects the accuracy of pathway-based time and phase prediction.

A practical obstacle in pathway-level metatranscriptomic time-series is that zero pathway values are ambiguous: they may reflect low abundance or limited detection, but may also result from biological absence, or the functional annotation procedure. Treating every zero as an exact quantitative observation may therefore impose unjustified certainty, and can bias trajectory estimation and downstream biomarker discovery (Chan and Li, 2024; Lyu et al., 2023). Gaussian processes (GPs) provide a principled Bayesian framework for modeling smooth time-series trajectories with uncertainty, but standard GP regression assumes fully observed responses and does not directly account for censoring (Basson et al., 2023). To address this limitation, we develop a censoring-aware GP (CAGP) framework tailored to metatranscriptomic time-series.

Longitudinal whole-microbiome metatranscriptomic datasets following food spoilage development and encoupled with sensory observations remain scarce. We therefore used a controlled experimental design to test, as a proof of concept, whether pathway-level transcriptional dynamics contain reproducible information about storage time and sensory spoilage progression. The present study was not designed to establish a universal spoilage signature or a routine shelf-life prediction tool. Rather, it evaluates the feasibility of longitudinal probabilistic modeling under a defined product, packaging atmosphere, and refrigeration range.

Specifically, this study addressed three questions within the investigated MAP broiler-meat system. First, do pathway-level metatranscriptomic profiles contain sufficient temporal structure to annotate the storage time of held-out samples? Second, is the inferred temporal structure associated with independently obtained sensory spoilage observations? Third, to what extent is the functional ordering learned at 4°C preserved in a separate time series collected at 6°C? We additionally explored which metabolic pathways and KEGG modules contributed most strongly to temporal and spoilage-phase discrimination. These analyses were intended to test the feasibility of the framework under controlled conditions rather than establish general applicability across food products or storage systems.

## 2. MATERIALS AND METHODS

### 2.1. Experimental dataset and pathway-level input data

This study used the longitudinal metatranscriptomic dataset generated and described in detail by Manninen et al., 2026. Briefly, fresh skinless broiler leg meat cuts were packaged under modified atmosphere and stored at two refrigeration temperatures, 4°C and 6°C. In the 4°C experiment, samples were collected daily from day 5 to day 14 after packaging, whereas in the 6°C experiment samples were collected daily from day 6 to day 10, reflecting the faster spoilage progression at the higher temperature. At each sampling day, three biological replicates with sufficient RNA yield were sequenced.

Sensory quality was evaluated as described previously, and the sensory scores were used to assign each sample to one of three spoilage phases: fresh product, early spoilage, or late spoilage. The phase assignments were therefore derived from sensory observations, not from storage day. In the 4°C dataset, days 5–10 were classified as fresh product, days 11–13 as early spoilage, and day 14 as late spoilage. In the 6°C dataset, days 6–7 were classified as fresh product, days 8–9 as early spoilage, and day 10 as late spoilage. These phase labels were used only for biological interpretation and evaluation of the model predictions; the Gaussian process models were fitted using chronological sampling day as the time variable.

The primary metatranscriptomic processing pipeline, including removal of ribosomal and host-derived reads, taxonomic profiling, functional profiling, and pathway quantification, was performed as described in Manninen et al., 2026. In brief, microbial reads were assigned to taxa and functionally profiled to obtain pathway-level activity estimates. Pathway abundances were normalized as transcripts per million (TPM), and the present analysis used log-transformed pathway activities, defined as log(TPM + 1), as input to the Gaussian process models.

For the 4°C dataset, pathway profiles were available for 30 samples corresponding to 10 sampling days with three biological replicates per day. For the 6°C dataset, pathway profiles were available for 15 samples corresponding to five sampling days with three biological replicates per day. The 4°C data were used as the main reference time-series for model fitting and leave-one-out validations, while the 6°C data were used to evaluate whether pathway dynamics learned at 4°C could transfer to an independent temperature condition.

### 2.2. Gaussian process time-series model

Gaussian processes (GPs) are a Bayesian non-parametric approach to regression that places a probability distribution over functions and yields predictive distributions for the inputs (Rasmussen and Williams, 2006). In time-series contexts, GPs can model smooth temporal trajectories, perform uncertainty quantification, and offer a principled way to perform interpolation and extrapolation from the data (Roberts et al., 2013).

A GP model *f*(*x*) is defined by a mean function *m*(*x*) and a covariance function *k*(*x, x*^′^). It is formally written as follows:

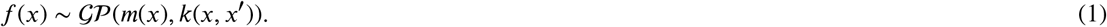

The kernel function is the most important part of a GP, as it encapsulates the assumptions we make about the latent function, such as its smoothness. The exponentiated quadratic kernel function is the most widely used for modeling (Rasmussen and Williams, 2006) with GPs and is defined as follows:

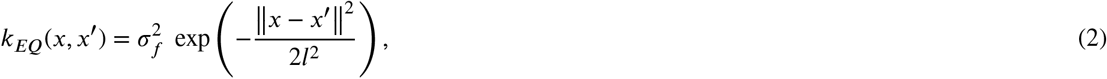

where *l* and 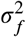 are the length-scale and variance. The length scale controls how the function oscillates, shaping the extent to which it fluctuates. In general, the model can only make reliable predictions within *l* units of the training data, meaning its ability to extrapolate beyond that point is inherently limited. The variance captures how far the function typically strays from its mean value. In GP modeling, we often assume that observations are noisy versions of the true function:

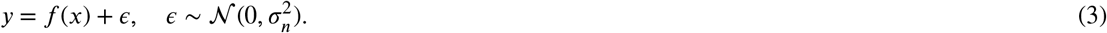

>Thus, when performing posterior inference with GPs there is a third parameter that we took into consideration: Gaussian noise 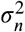, which represents the observation noise of the data in a GP.

The posterior over the latent function will be:

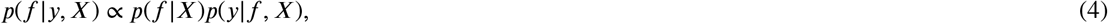

>where *p*(*f*|*X*) = *N*(*f*|*m, K*) is the GP prior over *f* (i.e., Equation (1)), and 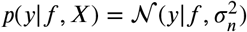 is the likelihood function (i.e., Equation (3)). For a set of outputs *y*_1_, …, *y*_*N*_ the GP likelihood function will then be:

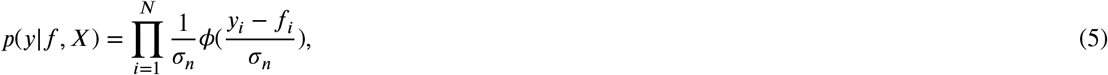

>where *ϕ*(*⋅*) is the standard normal probability density function.

When fitting a GP with a given set of input *t* and output *y* we aimed to find the optimal hyperparameters 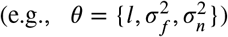 that maximize the marginal likelihood of the model:

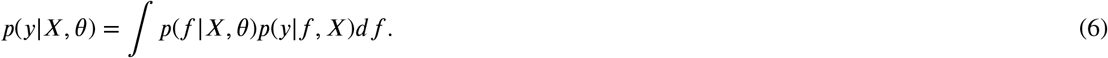

### 2.3. GP for censored data

Most GP tutorials assume that observations are directly measured (i.e., Gaussian noise around latent values), but in many cases, the apparent sparsity of the data may be partly attributable to value censoring, for example when measurements fall below the instrument’s limit of detection (LOD) rather than being truly absent (Gorm Hoffmann et al., 2025).

In this case, the data will be composed of many observed zero values, which are not real zeros but merely small values which appear as zeros due to the LOD.

To accommodate this, we modified the likelihood function to reflect the censoring mechanism, which is a standard approach for addressing the issues of censored data as described in many studies (Gammelli et al., 2020; Gorm Hoffmann et al., 2025; Groot and Lucas, 2012; Karlova et al., 2024). Let *y*_*i*_ denote the observed value for observation *i*, and let *τ* denote the LOD threshold. The observations can be partitioned in two sets: uncensored observations, where *y*_*i*_ > 0, and censored observations, where *y*_*i*_ = 0. For the uncensored observations, the true value is known, and the standard Gaussian likelihood applies:

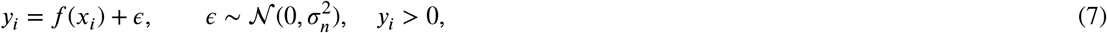

>and

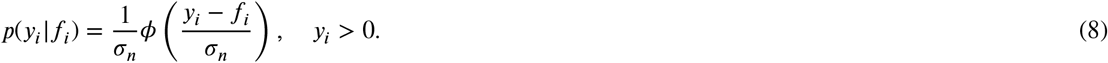

>For the censored observations, we do not observe the true value; we only know that it fell below *τ*. In this case, we considered an additional noise for the observations:

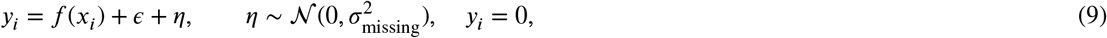

>and the appropriate likelihood is the probability that the latent function plus noise lies below the threshold:

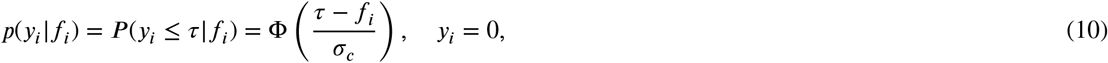

>where 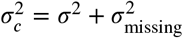, and Φ(*⋅*) is the standard normal cumulative distribution function (CDF), also referred to as the probit function. This is a natural choice, as it returns the probability that a Gaussian random variable with mean *f*_*i*_ and variance 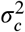 takes a value at or below the detection threshold. Our introduction of 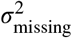 turns this into a soft threshold, with the normal CDF around *τ* denoting the probability of censoring.

The combined likelihood function over all *N* observations is therefore:

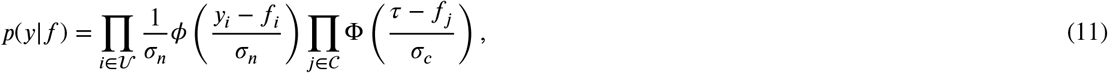

>where *U* and *C* denote the index sets of uncensored and censored observations, respectively.

Because the likelihood in Equation (11) is no longer Gaussian due to the probit terms, the posterior *p*(*f*|*y, X*) is no longer analytically tractable. Hence, we resorted to approximate inference using Expectation Propagation (EP) to approximate it. We provide a Python implementation of the censoring-aware GP (CAGP) on GitHub. Our implementation was based on (Gammelli et al., 2020), which is built on top of GPy (GPy, last accessed date 03/08/2025), with the necessary modifications to turn *τ* into a lower LOD instead of an upper LOD, and the addition of the extra noise *σ*_missing_ that turns it into a soft threshold. Previous work assumes a hard threshold: anything below *τ* is censored with a 100% probability, anything above is not.

### 2.4. Time point annotation of metatranscriptomic samples

Time annotation of the metatranscriptomic samples was performed following the Bayesian GP-based approach of Nushi et al., 2025, adapted here to account for the censored nature of the data. Specifically, a separate censored GP model *f*_*p*_(*x*) was fitted to the reference pathway activity time-series for each pathway *p*, using the exponentiated quadratic kernel as described in Section 2.2. Given a target metatranscriptomic sample *S* with pathway activity values {*a*_1,*S*_, *a*_2,*S*_, …, *a*_*P,S*_}, and fitted censored GP models *M*_1_, *M*_2_, …, *M*_*P*_ from the reference time-series, the likelihood of the sample having been collected at time *t* was computed by accounting for the censoring status of each pathway activity value. Let *U* and *C* denote the index sets of uncensored and censored pathway activity values in sample *S*, respectively. The likelihood of the sample at time *t* is then:

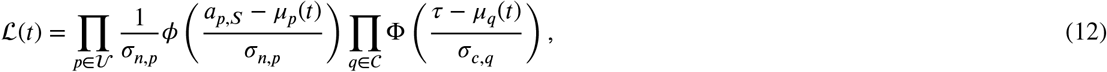

>where *µ*_*p*_(*t*) and *µ*_*q*_(*t*) are the posterior predictive means of *M*_*p*_ and *M*_*q*_ at time *t, ϕ*(*⋅*) is the standard normal probability density function, Φ(*⋅*) is the standard normal cumulative distribution function, *τ* is the limit of detection threshold, 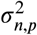 is the model specific noise for uncensored data, and 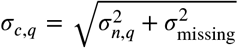 is the model specific noise for censored data. The first product in Equation (12) evaluates the Gaussian likelihood of each uncensored pathway activity value under the GP posterior at time *t*, while the second product evaluates the probability that each censored pathway activity value falls below the detection threshold *τ*, again under the GP posterior at time *t*.

The time annotation was then obtained by maximizing the sum of log-likelihoods over a discrete time grid {*t*_1_, *t*_2_, …, *t*_*T*_ }:

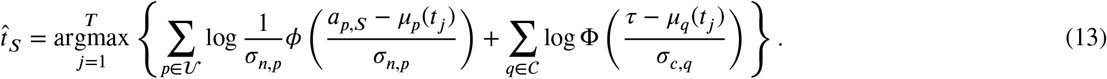

>Pathways whose activity values in the target samples were better described by a time-invariant model than by their corresponding censored GP model were excluded from the annotation, following the quality control procedure described in (Nushi et al., 2025). The key difference from the original formulation of (Nushi et al., 2025) is two fold: first, rather than modeling individual gene expression profiles, each GP here captures the temporal dynamics of a pathway activity score, aggregating the transcriptional signal of functionally related genes into a single time-indexed quantity; second, the likelihood used for time annotation explicitly accounts for censoring, replacing the standard Gaussian likelihood with a mixed likelihood that uses a Gaussian density for uncensored observations and a probit term for the censored observations (i.e., zero-valued observations).

### 2.5. Selection of the detection limit *τ* and *σ*_missing_ for left-censored zero observations

#### 2.5.1. Data scale and motivation

All pathway abundances were analyzed on the log scale

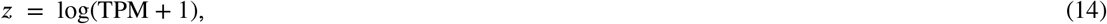

>so that *z* ≥ 0, and *z* = 0 corresponds exactly to TPM = 0. The dataset contains replicate measurements for each pathway at each discrete time point *t ∈* {day 5, …, day 14} (columns *tA, tB, tC*), where replicates are independent within a time point and replicate labels are not tracked across time.

Although *z* = 0 is a valid transformed value, we treat zero observations as unreliably quantified in the sense that they carry limited information about the underlying pathway abundance beyond being “low”. Accordingly, we model zeros with a left-censoring mechanism: a zero observation is assumed to indicate that the latent log-abundance lies below a threshold *τ*, rather than being an exact point measurement. The threshold *τ* determines how informative zeros are in the likelihood and therefore affects time annotation performance. We select *τ*, and *σ*_missing_ in a data-driven manner, using replicate information and probit model.

#### 2.5.2. Replicate-based detectability model

To determine the censoring threshold *τ*, and *σ*_missing_ for the censoring-aware Gaussian process (CAGP) model, we first modeled the probability that an observation is recorded as zero as a function of abundance on the log(TPM + 1) scale. For each pathway *p* and time point *t*, let *z*_*pt*1_, …, *z*_*ptR*_ denote the replicate measurements, and let

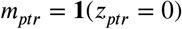

>indicate whether replicate *r* was observed as zero. We summarized the replicate values by their highest value,

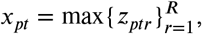

>and assigned this same proxy value *x*_*pt*_ to all replicates in the set. Here, we make the implicit assumptions that within a pathway-time group, the highest observed replicate is the best available proxy for latent abundance, as it has the strongest signal, while lower or zero replicates are partly due to detection failure.

A probit regression model was then fitted to these replicate-level data:

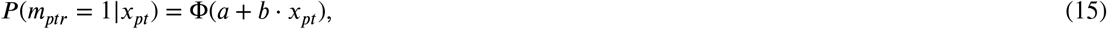

>where Φ(*⋅*) is the standard normal CDF and *a, b* are the intercept and slope estimated by maximum likelihood. This model can be parameterized in terms of the censoring threshold and noise: setting *a* = *τ*/*σ*_missing_ and *b* = ™1/*σ*_missing_, the probit becomes

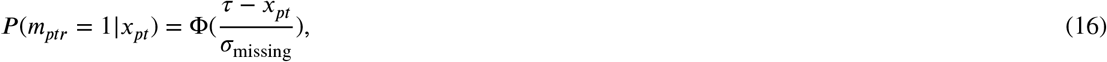

>which is the probability that a Gaussian random variable with mean *x*_*pt*_ and standard deviation *σ*_missing_ falls below *τ*.

## 3. RESULTS

### 3.1. Data description and preprocessing

Our data consisted of metatranscriptomic samples collected from packaged chicken products in consecutive days. 3 samples (i.e., biological replicates) were collected every day, each from a different package, which was removed after the sampling. The same procedure was conducted in 4°C and 6°C. In total 30 samples were collected during 10 consecutive days (day 5 - day 14) in the 4°C experiment which we label {*t*A, *t*B, *t*C} (*t ∈* {5, …, 14}) to differentiate samples by day and replicate. Similarly 15 samples were collected during 5 consecutive days in the 6°C experiments which we label {*t*D, *t*E, *t*F} (*t ∈* {6, …, 10}) accordingly.

We removed bacteria whose fraction of total reads were less then 5% and considered only those pathways which were associated with at least one bacterium. 938 metabolic pathways were identified in the 4°C samples which were associated with 9 different strains of bacteria: *Carnobacterium divergens, Carnobacterium maltaromaticum, Pseudolactococcus piscium, Leuconostoc gelidum/gasicomitatum, Latilactobacillus sakei, Pseudomonas lundensis, Vagococcus proximus*, and *Yersinia intermedia*. 284 metabolic pathways were identified in the 6°C samples, which were associated with 3 different strains of bacteria: *Carnobacterium divergens, Carnobacterium maltaromaticum*, and *Vagococcus proximus*. There were 283 metabolic pathways in common between the samples of 4°C and 6°C.

In addition to chronological sampling day, samples were assigned to spoilage phases based on sensory scores, yielding fresh, early spoilage, and late spoilage groups as described in Section 2.1. This phase-level labeling was used as biologically interpretable feature for evaluating transfer across temperatures. The phase labels provide an important complement to exact day prediction because different storage temperatures are expected to compress or expand the temporal trajectory while preserving broad spoilage states.

To visualize the global structure of the dataset, we performed principal component analysis on the shared pathway abundance matrix after log-transformation (i.e., log(TPM+1)), as shown in Figure 1. The projection showed an ordered progression of the samples over storage time and indicated that the 4°C and 6°C datasets were related but not identical in their global transcriptional organization. This is consistent with the expectation that temperature alters the rate of spoilage progression without fully erasing the underlying functional succession pattern. For this reason, we treated the 6°C series as a shifted and (potentially non-linearly) compressed but biologically comparable target domain, where information from the temporal model trained on 4°C samples can be transferred.

**Figure 1.**
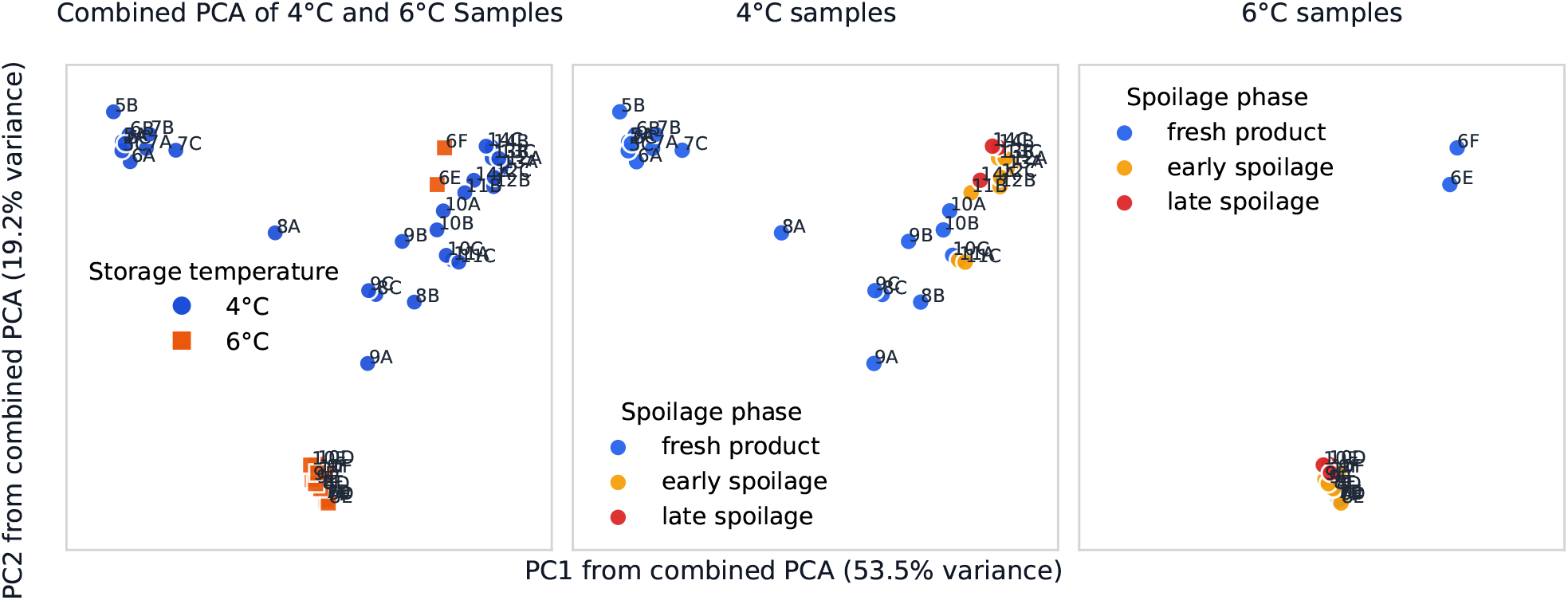
Principal component analysis (PCA) of pathway activity profiles across storage conditions. Left: PCA of shared pathways shows a clear separation between 4°C (blue) and 6°C (orange) samples along the second principal component (PC2), indicating a strong temperature-driven effect. Middle and right: PCA of shared pathways within each temperature condition, colored by spoilage phase (fresh, early, late), show a consistent temporal progression from fresh to late spoilage. While the progression is gradual at 4°C, it appears more separated at 6°C, suggesting that temperature primarily modulates the rate of biological change.

### 3.2. Cross-validation analyses on 4°C samples

#### 3.2.1. Leave-one-replicate-out analysis on 4°C samples

We first assessed whether the censoring-aware GP (CAGP) model could recover sampling time within the 4°C experiment using a leave-one-replicate-out (LORO) design. For evaluating the limit of detection *τ* and *σ*_missing_, we followed the procedure described in Section 2.5 and obtained *τ* = 3.94, *σ*_missing_ = 1.83. In the Supplementary Material File 3 section S.1 we include a sensitivity analysis for different values of (*τ, σ*_missing_). For each held-out sample, the model combined pathway-level likelihoods across the fitted GP trajectories and assigned the time point *t* with the highest posterior, following Equation (13), from the uniformly spaced time grid *t ∈* [5.0, 5.1, 5.2, …, 14.0] (the earliest samples are from day 5 and the latest samples are from day 14;thus the time grid spans from day 5 to day 14). This internal validation setting tests whether the pathway trajectories are reproducible enough across biological replicates to support precise temporal annotation rather than merely broad phase discrimination.

The model predicted the sampling day with high accuracy, with a mean absolute error (MAE) of approximately 0.43 days from the true day across all the samples. We compared CAGP with the normal GP modeling, where data censoring is not taken into consideration, and we obtained MAE=1.31 days. Figure 2 shows the MAE of the replicate predictions at each time point for CAGP and normal GP. Most held-out samples were assigned either to the correct day or to an immediately neighboring day, indicating that the fitted pathway trajectories captured the temporal progression of spoilage at a fine resolution. Figure 3 shows the accuracy of the models (CAGP and normal GP), measured as the proportion of the samples whose predictions in the LORO are within a certain time tolerance of their true time.

**Figure 2.**
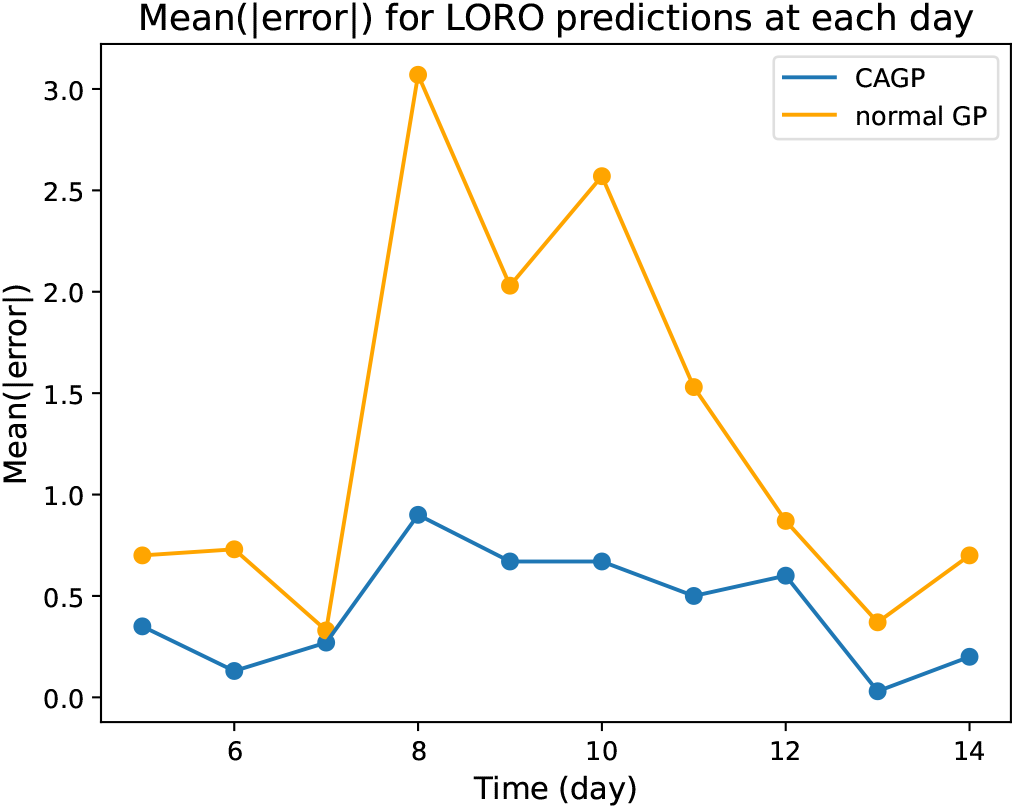
Mean absolute error of the replicates predicted at each time point, using LORO cross-validation on the 4°C samples. Mean(|error|) is evaluated as the absolute value of the difference between the predicted time 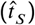 and the true time (*t*_*S*_) of the sample 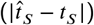.

**Figure 3.**
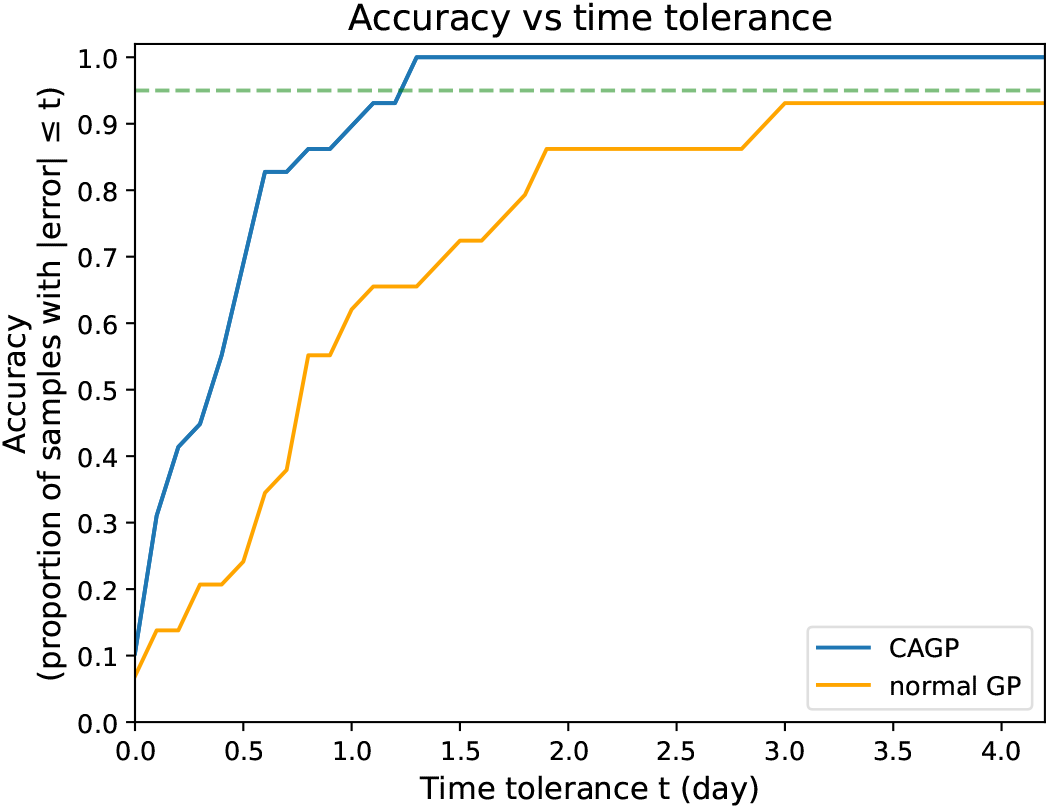
Accuracy of the CAGP and GP for predicting the time point on the LORO 4°C samples within a given time tolerance from their true time point. The curve shows the fraction of samples whose absolute prediction error is less than or equal to a tolerance *t*. Prediction accuracy increases with time tolerance, with approximately 95% within 1.2 days (green line) for CAGP.

Then we mapped each time prediction to a spoilage phase according to the schema mentioned in Section 2.1 (i.e., days 5–10.4: fresh product, days 10.5–13.4: early spoilage, days 13.5–14: late spoilage). As shown in Figure 4 (left-panel), at the phase level, the predictions were concordant with the sensory labels (except for a couple of late samples on the upper edge of fresh product phase), showing that the latent temporal signal learned from pathway activity was aligned with the biologically relevant progression from fresh to spoiled meat.

**Figure 4.**
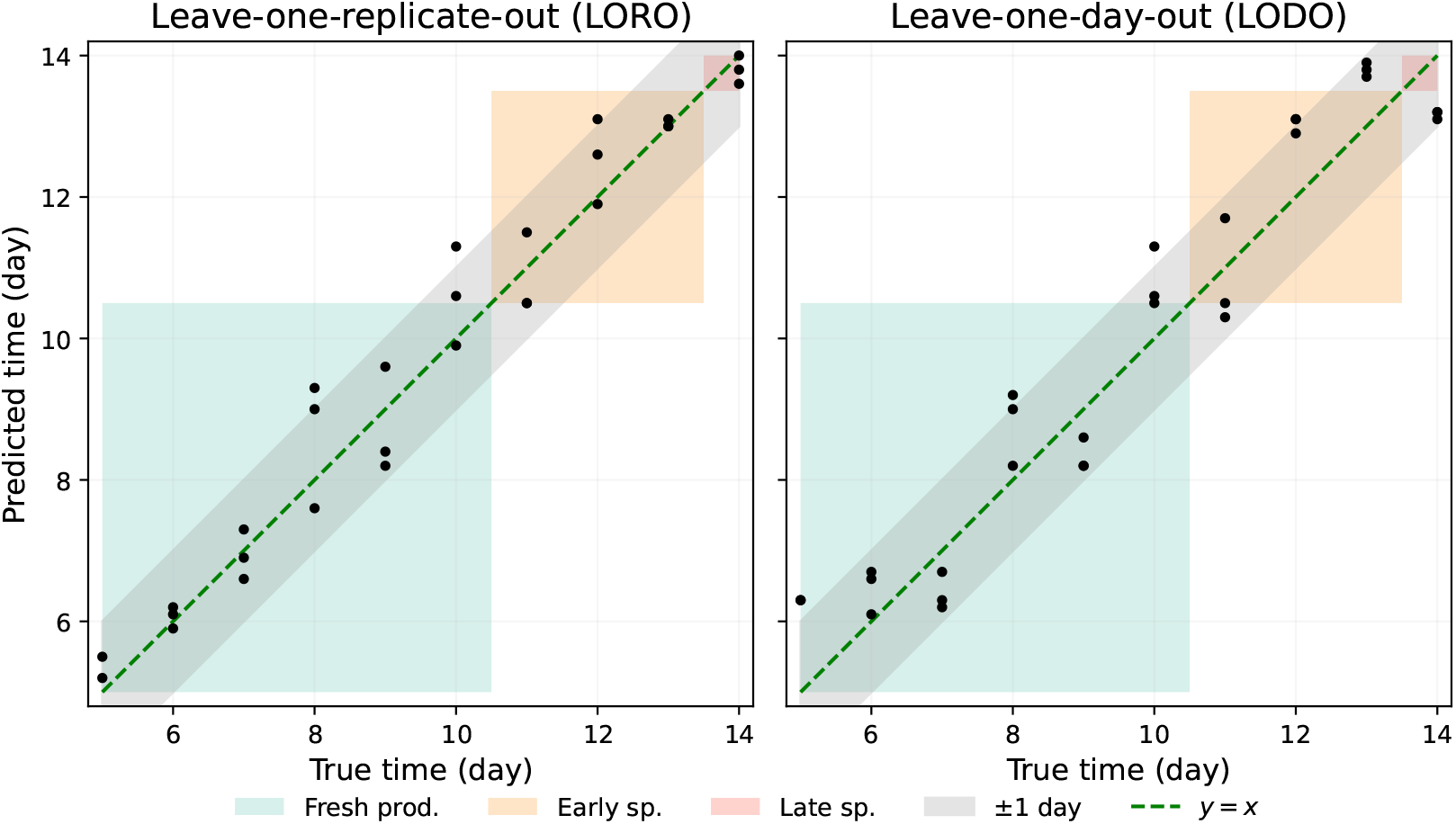
Comparison of LORO and LODO storage-time annotation at 4°C using the CAGP. In LORO validation (left), one biological sample was excluded while the other samples collected on the same storage day remained in the reference dataset. In LODO validation (right), all three biological samples from the target storage day were excluded simultaneously. Points represent individual samples, the dashed green line indicates perfect correspondence between predicted and true storage day, and the gray region indicates predictions within ±1 day of the true storage day. The MAE was 0.43 days for LORO and 0.78 days for LODO. The colored diagonal regions indicate combinations in which true and predicted storage days fall within the same temporal intervals corresponding to the sensory-derived phases observed in this dataset; the phase labels were not used to train the model.

These results show that pathway-level metatranscriptomic patterns were reproducible across biological replicates collected on the same storage day. The more stringent question of whether the model could annotate samples from an entirely unseen storage day was evaluated using leave-one-day-out validation (LODO).

#### 3.2.2. Leave-one-day-out analysis on 4°C samples

We next evaluated the CAGP using leave-one-day-out (LODO) cross-validation. In each validation fold, all three biological replicates from one storage day were excluded, leaving no samples from the target day in the reference time series. The excluded samples were then annotated using pathway trajectories fitted to the remaining nine storage days. Under this more stringent validation design, the CAGP predicted storage time with a mean absolute error of 0.78 days. Although the error was higher than the 0.43-day MAE obtained in the LORO analysis, the average prediction error remained below the one-day sampling interval. This indicates that the pathway trajectories contained sufficient temporal structure to support interpolation across a completely unobserved storage day.

The increase in error relative to LORO validation was expected because the model could no longer use biological replicates collected on the target day. Instead, it had to infer the missing temporal position from pathway trajectories learned from the surrounding days. For the first and last storage days, the analysis additionally required extrapolation because observations were available on only one side of the excluded time point. Figure 4 (right-panel) shows the alignment between the true time and the predicted time for each sample in the LODO cross-validation.

Phase-level agreement was lower in LODO validation than in LORO validation. The disagreements were con-centrated among samples located near the transitions between spoilage phases, as illustrated by Figure 4 (right-panel). Because predicted phases were obtained by applying fixed boundaries to the continuous time estimates, relatively small temporal errors could change the assigned phase. For example, a sample from day 11 predicted slightly before day 10.5 was classified as fresh rather than early spoilage, despite an error of less than one day. Similarly, predictions around the 13.5-day boundary could switch between early and late spoilage. Thus, the categorical discrepancies were between adjacent phases rather than between biologically distant spoilage states.

### 3.3. Cross-temperature annotation of 6°C samples using the 4°C series as the reference

We then evaluated whether pathway trajectories learned from the 4°C time-series could be used to annotate samples collected from an independent chicken spoilage experiment stored at 6°C. Because elevated temperatures are expected to accelerate the spoilage process, exact alignment of the predicted day and the observed day was not expected. Instead, the main question was whether the temporal ordering and spoilage phase progression remained recoverable under the temperature shift.

We evaluated the limit of detection *τ* and *σ*_missing_, using the 4°C data as described in Section 2.5. Then, we used the 4°C time-series replicates to fit the data censoring-aware GP (CAGP) models as shown in Section 2.4. The transferred model assigned later 6°C samples progressively later positions on the 4°C reference timeline as shown in Figure 5, demonstrating that the inferred pathway programs retained interpretable temporal information across temperatures. As expected under the higher storage temperature, the transferred model did not preserve a one-to-one correspondence between chronological sampling days at 6°C and the 4°C reference timeline. Instead, the 6°C samples were systematically displaced toward later positions on the 4°C trajectory. Spearman’s correlation was used because exact numerical correspondence between storage days at the two temperatures was not expected and because several predictions reached the upper boundary of the reference interval, and it yielded a *ρ* = 0.95. Day 6 samples mapped near the late-fresh boundary of the 4°C series; Day 7 mapped predominantly to early spoilage phase, day 8 samples mapped to late-early spoilage and late spoilage positions, and day 9–10 samples saturated at the last 4°C time point. Thus, the cross-temperature prediction should not be interpreted as exact calendar-day transfer. Rather, it shows that the model recognized the accelerated progression of the 6°C experiment and placed these samples along the expected direction of spoilage advancement. The compression of day 8–10 samples into the late region is consistent with rapid convergence toward a spoiled functional state at elevated temperature, it also shows a possible upper bound effect as predictions reached the upper bound of the reference interval which is day 14.

**Figure 5.**
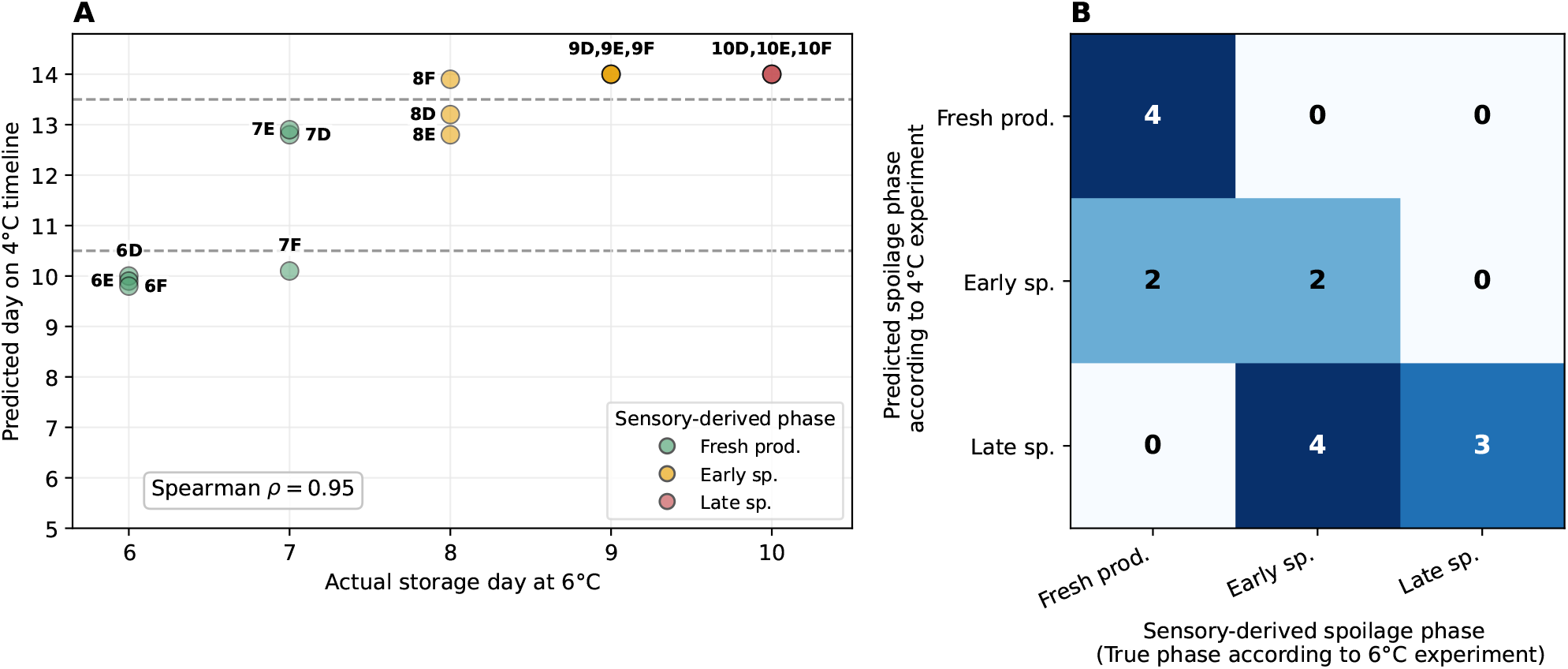
Quantitative evaluation of cross-temperature annotation of the 6°C samples using the 4°C time series as the reference. (A) Actual storage day at 6°C versus the equivalent position predicted by the CAGP on the 4°C reference timeline. Each point represents an individual sample, with sample identifiers shown next to the points and colors indicating the original sensory-derived spoilage phases. Increased opacity indicates overlapping or closely positioned observations. Horizontal dashed lines mark the boundaries used to convert predicted equivalent days into fresh (<10.5 days), early-spoilage (10.5 to <13.5 days), and late-spoilage (≥13.5 days) phases. Spearman’s correlation coefficient quantifies the preservation of temporal ordering across temperatures. Predictions reaching day 14 correspond to the upper limit of the available 4°C reference interval and should therefore be interpreted as boundary-limited estimates rather than precise equivalent days. (B) Confusion matrix comparing the sensory-derived phases of the 6°C samples (columns) with the phases inferred from their predicted positions on the 4°C reference timeline (rows)

These findings support the central claim of the study: although storage temperature changes the pace of spoilage, a shared functional metatranscriptomic signature of spoilage progression is preserved across conditions. The cross-temperature analysis suggests that the model is capturing biologically meaningful spoilage dynamics rather than overfitting temperature-specific noise in the 4°C dataset.

### 3.4. Cross-entropy ranking and cross-temperature overlap of spoilage-informative pathways

We next asked which pathway–taxon trajectories carried the strongest information about storage time and about sensory spoilage phase. For this analysis, the 4°C samples were grouped as fresh product on days 5–10, early spoilage on days 11–13, and late spoilage on day 14. The 6°C samples were grouped as fresh product on days 6–7, early spoilage on days 8–9, and late spoilage on day 10. This phase mapping allowed us to distinguish markers of chronological sampling day from markers of the biological spoilage state, which is important because the 6°C experiment is expected to progress faster than the 4°C reference time-series.

We ranked pathways using the cross-entropy score (i.e., Equation 17), where lower values indicate that the pathway assigns high probability to the target day while assigning lower probability to other days.

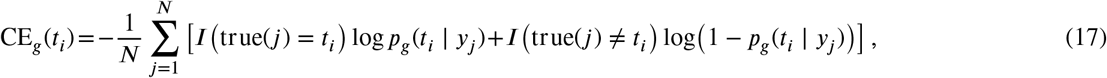

>where *g* represents a specific pathway, *N* is the number of observations in that pathway, *t*_*i*_ is the time point for which we are measuring the cross-entropy in pathway *g*, and true(*j*) is the true time label of sample *j*. The files with the pathways’ cross-entropy scores can be found in the supplementary material as specified in Section 5.

To focus the ranking on observed pathway activity, we considered only pathway–day entries with at least one observation at the target day. This criterion avoids selecting pathways whose apparent time specificity is driven only by non-detection at the target time point.

The observed cross-entropy ranking retained cross-temperature agreement, particularly after collapsing species-resolved pathway–taxon entries to KEGG module identifiers. Among the top 50 observed cross-entropy markers per spoilage phase, exact pathway–taxon overlap between 4°C and 6°C was 9 markers in fresh product, 11 in early spoilage, and 16 in late spoilage, whereas module-level overlap was 15, 15, and 26 modules, respectively, as shown in Figure 6.

**Figure 6.**
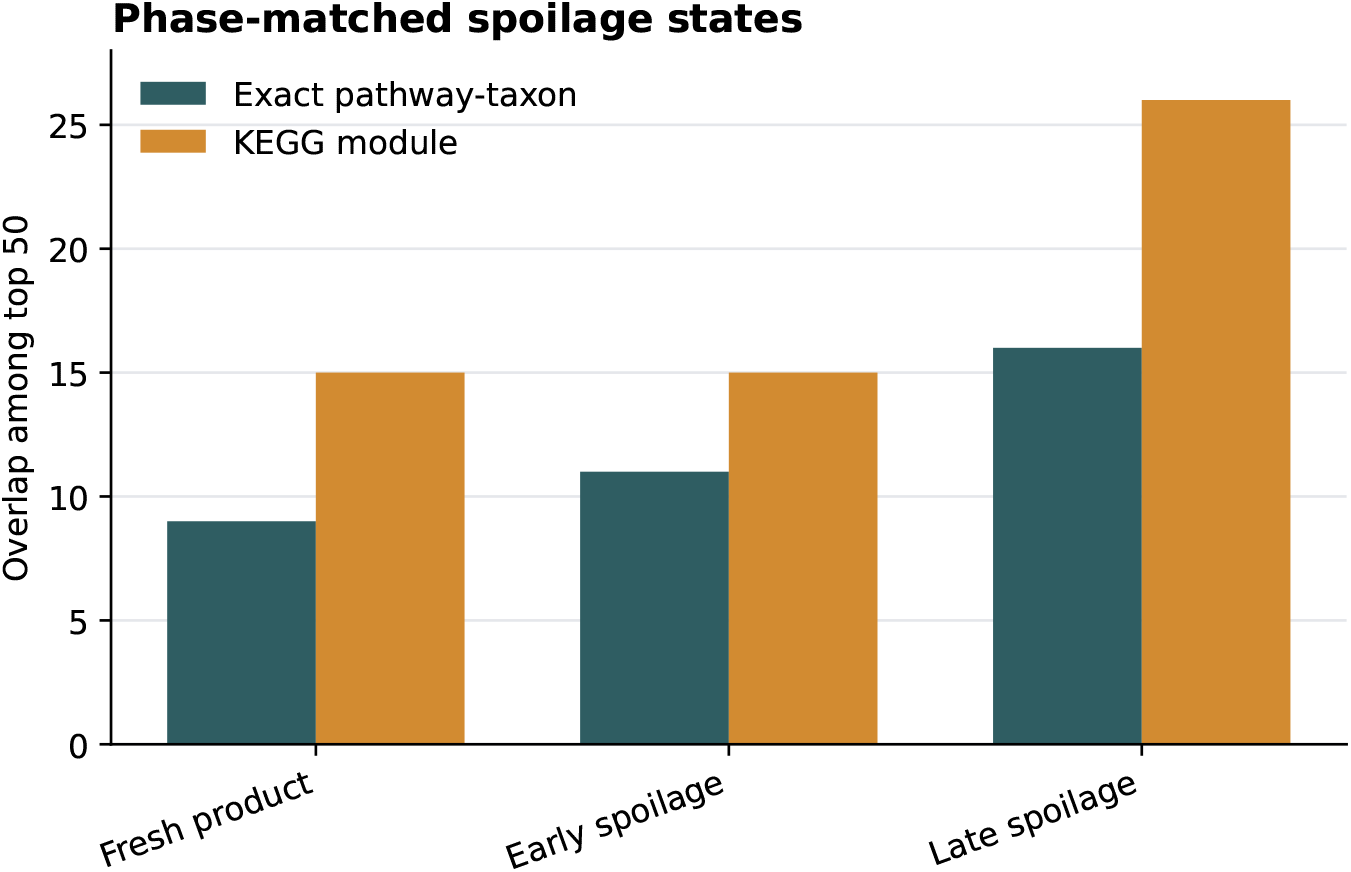
Cross-temperature overlap among observed cross-entropy-ranked pathway markers after grouping days into spoilage phases. Pathway–day entries with no observations at the target day were excluded before ranking. Bars show the number of shared markers among the top 50 cross-entropy-ranked entries when requiring exact pathway–taxon matches or when collapsing entries to KEGG module identifiers.

To determine whether the overlap between the 4°C and 6°C rankings exceeded that expected by chance, we performed a permutation analysis using 100,000 random selections from the eligible pathway sets. Among the top 50 entries, no significant enrichment was detected for fresh product. Early spoilage showed suggestive pathway–taxon enrichment, but this did not remain significant after multiple-testing correction. In contrast, late spoilage showed strong cross-temperature enrichment. The observed pathway–taxon overlap was 16 compared with 6.20 expected by chance, corresponding to a 2.58-fold enrichment (*p*_adj._ = 0.00036). At the KEGG-module level, 26 shared modules were observed compared with 15.21 expected, corresponding to a 1.71-fold enrichment (*p*_adj._ = 0.00036). Cross-entropy produces a continuous ranking with no natural cutoff separating informative from non-informative pathways. We therefore used the top 50 entries from each temperature as a pragmatic primary comparison. Using lists of equal length prevents differences in list size from influencing the observed overlap. Because the choice of 50 was not biologically or statistically unique, we repeated the analysis using the top 25 and top 100 entries to assess cutoff sensitivity. Sensitivity analyses using the top 25 and top 100 entries confirmed that late-spoilage pathway–taxon overlap remained significantly enriched across all three cutoffs, whereas results for fresh and early spoilage were more cutoff-dependent (Supplementary Material File 3 section S.2). These findings indicate that cross-temperature enrichment of the highest-ranked pathway markers was most consistent during late spoilage. Overlap during fresh product and early spoilage was more dependent on the ranking cutoff and level of functional resolution.

We therefore focused the subsequent biological interpretation on the 26 KEGG modules shared between the late-spoilage rankings. The shared late-spoilage modules included functions associated with carbohydrate metabolism, amino-acid metabolism, energy metabolism, nucleotide metabolism, and cofactor and vitamin metabolism (Figure 7). Carbohydrate-metabolism modules may reflect the utilization and fermentation of available substrates during microbial growth. Amino-acid-metabolism modules are particularly relevant to spoilage because amino-acid degradation can produce amines, sulfur-containing compounds, aldehydes, and other metabolites associated with undesirable odors. Energy-metabolism and cofactor-related modules may reflect the redox and biosynthetic requirements of microorgan-isms growing under refrigerated modified-atmosphere conditions. Nucleotide-metabolism modules are more likely to reflect microbial proliferation and changing biosynthetic demands than to contribute directly to off-odor formation. These functional associations are biologically plausible, but the cross-entropy analysis does not establish that the identified modules directly cause sensory deterioration.

**Figure 7.**
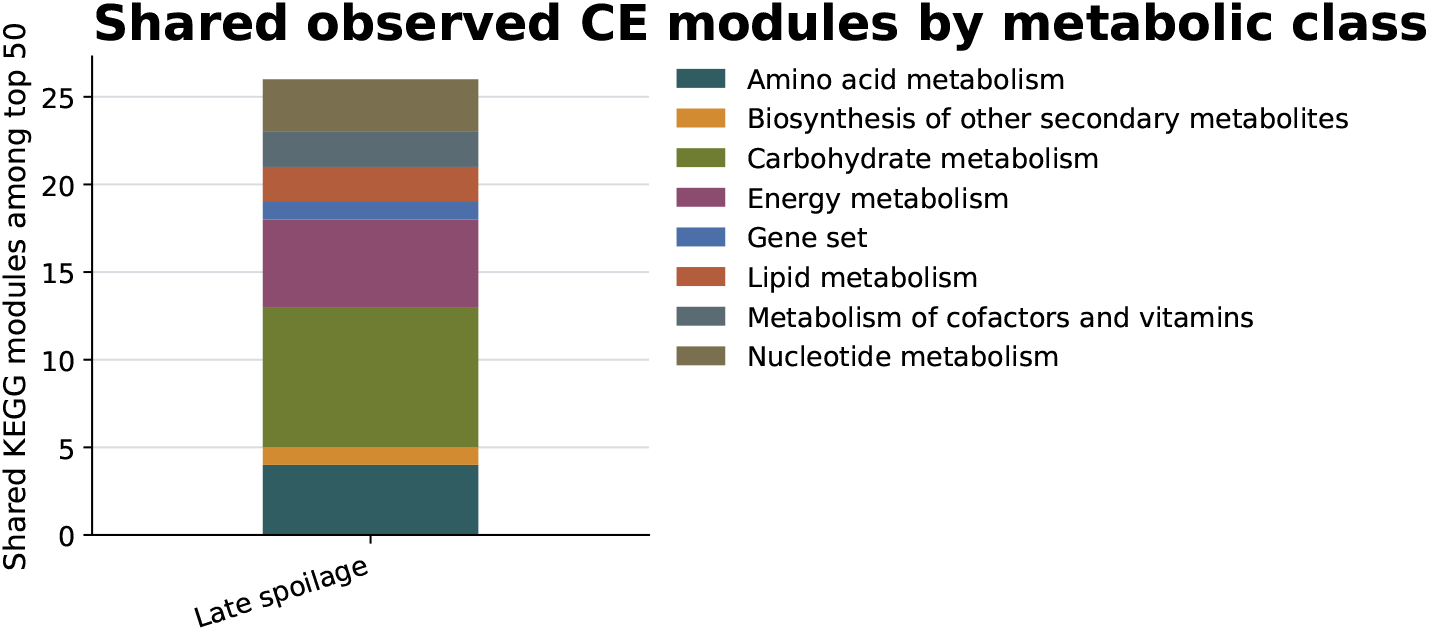
Metabolic classification of the shared late-spoilage KEGG modules. The stacked bar shows the metabolic classes represented among the 26 KEGG modules shared between the top 50 cross-entropy-ranked entries from the 4°C and 6°C datasets during late spoilage. Permutation analysis showed that this overlap was significantly greater than expected by chance (expected overlap = 15.21 modules; fold enrichment = 1.71; empirical (p<0.0001); Benjamini–Hochberg-adjusted p = 0.00036).

## 4. DISCUSSION AND CONCLUSION

The sensory-defined spoilage phases and the descriptive analysis of microbial and pathway activity in these samples were reported by Manninen et al., 2026. Here, we reanalyze the longitudinal pathway profiles to estimate continuous trajectories, predict storage time, assess agreement with sensory phases, and test cross-temperature transfer and pathway-marker overlap. Peer-reviewed studies of refrigerated chicken independently show that microbial communities and sensory quality change during storage, while temperature affects the rate of deterioration (Balamatsia et al., 2006; Yimenu et al., 2019).

Our first research question was whether pathway-level metatranscriptomic profiles can be modeled to predict the storage time of individual samples. The results supported this. In the 4°C experiment, the censoring-aware Gaussian process model recovered sampling time with high accuracy in LORO and LODO validation, with a mean absolute error of approximately 0.43 and 0.78 days, respectively. Most predictions were either correct or close to the true sampling day, showing that pathway activity contained a reproducible temporal signal across biological replicates.

Also, regarding our second research question, whether the same pathway-based model could distinguish sensory-defined spoilage phases, the answer was positive. Although the model was trained using chronological sampling days rather than sensory phase labels, its predicted times were largely concordant with the fresh, early-spoilage, and late-spoilage classifications in the LORO cross-validation. This result is important because it shows that the temporal signal learned from pathway activity was not only a mathematical ordering of samples but also reflected biologically meaningful spoilage progression. In other words, the model recovered a functional trajectory that aligned with the sensory deterioration of the product.

Our third research question was whether storage temperature affects the ability of the model to predict storage day and spoilage phase. The cross-temperature analysis showed that exact calendar-day prediction did not transfer directly from 4°C to 6°C, as expected, because the higher temperature accelerated spoilage progression. However, the 6°C samples were still placed along the expected direction of the 4°C spoilage trajectory, with later 6°C samples mapping to later positions in the 4°C reference timeline. This suggests that temperature changes the rate of spoilage but does not completely change the underlying functional progression. Therefore, the model should not be interpreted as providing a one-to-one conversion between days at different temperatures, but rather as capturing selected aspects of the functional progression of spoilage that are preserved across refrigeration conditions within the investigated experimental system.

The cross-entropy analysis qualifies, rather than contradicts, the cross-temperature prediction results. The GP-based annotation combines likelihood information across the complete set of modeled pathways, whereas the overlap analysis applies a more stringent criterion by asking whether identical pathway–taxon entries or KEGG modules occur among the highest-ranked markers at both temperatures. Consequently, successful temporal annotation does not require the same individual pathways to occupy the highest ranks in both datasets. Predictive information may be distributed across several correlated or functionally related pathways whose relative importance changes with temperature. The limited enrichment of top-ranked markers during the fresh and early-spoilage phases therefore does not imply an absence of shared temporal information during these phases. Instead, it suggests that this information may be less concentrated in a common set of dominant markers. By contrast, the consistent enrichment observed during late spoilage indicates greater convergence of the two temperature conditions toward a common set of highly informative functional processes as spoilage advances. The 26 KEGG modules shared between the late-spoilage rankings included functions associated with carbohydrate metabolism, amino-acid metabolism, energy metabolism, nucleotide metabolism, and cofactor and vitamin metabolism. This composition is biologically plausible for advanced microbial spoilage. Carbohydrate-metabolism modules may reflect the utilization and fermentation of available substrates, while amino-acid degradation can generate amines, sulfur-containing compounds, aldehydes, and other metabolites associated with undesirable odors. Energy-metabolism and cofactor-related modules may reflect the redox and biosynthetic requirements of microorganisms growing under refrigerated modified-atmosphere conditions. Nucleotide-metabolism modules may primarily indicate microbial proliferation and changing biosynthetic demands rather than directly contributing to off-odor formation. These interpretations are consistent with peer-reviewed evidence linking microbial use of meat nutrients to spoilage-related volatile compounds (Casaburi et al., 2014). The present analysis identifies associations between pathway activity and spoilage progression; it does not establish which pathways produce the sensory changes. Nevertheless, the recurrence of these modules should be interpreted as an association with the late-spoilage state rather than evidence that they directly cause sensory deterioration. Establishing such mechanistic relationships would require integration with metabolomic, volatile-compound, and individual sensory-score measurements.

In the broader context, this work continues two lines of previous research. First, it extends the metatranscriptomic study of poultry meat spoilage in (Manninen et al., 2026), moving from phase-wise comparison of pathway activity to continuous time-series modeling of pathway trajectories. Second, it adapts the supervised Bayesian time-annotation framework of (Nushi et al., 2025) from transcriptomic time-series analysis to pathway-level metatranscriptomic data. The main methodological contribution is the incorporation of censoring-aware likelihoods into the time-annotation framework, which is important because zeros in metatranscriptomic pathway profiles may reflect non-detection rather than true biological absence. By treating zero values as censored observations, the model better reflects the measurement process and avoids assigning excessive confidence to uncertain low-abundance signals.

These results also have practical implications for shelf-life research. Traditional microbiological approaches such as culturing, PCR, and amplicon sequencing are valuable, but they mainly describe microbial presence, abundance, or species composition in a spoilage microbiome. They do not always explain when sensory spoilage becomes detectable, especially in cold-stored MAP meat where high bacterial counts may occur some days before the product is rejected. In addition, strain-related diversity challenges all spoilage microbiome analyses since it plays more important role than a species status (Casaburi et al., 2014; Illikoud et al., 2019; Pothakos et al., 2014; Sanders et al., 2015). The present study supports the idea that functional molecular data can provide an additional layer of information by describing what the microbial community is doing over time. In the long term, such functional signatures could contribute to more predictive shelf-life assessment, better identification of spoilage mechanisms, and improved monitoring of product quality.

However, some limitations should be considered. First, our study was based on a limited experimental setting: two broiler meat datasets, two storage temperatures, and a relatively small number of samples. Although the LORO results at 4°C were strong and the 6°C transfer analysis was encouraging, broader validation is needed across production lots, packaging conditions, meat types, and independent datasets. Second, the 6°C dataset covered a shorter and faster spoilage trajectory than the 4°C dataset, meaning that cross-temperature predictions were necessarily evaluated at the phase level rather than as exact day predictions. A larger multi-temperature design would be needed to explicitly model temperature as a continuous driver of spoilage rate.

The third limitation is that the current model treats pathways independently when combining likelihoods for time annotation. This simplifies inference and interpretation, but it ignores correlations between pathways, taxa, and metabolic modules. In reality, spoilage is driven by interacting microbial communities and coordinated metabolic processes. Future work could therefore explore multi-output Gaussian processes, hierarchical Bayesian models, or latent-factor approaches that model shared structure between pathways. Fourth, the censoring threshold and missingness parameters were estimated from replicate information using assumptions about detectability. Although this is a practical solution, the true detection process may vary across pathways, taxa, sequencing depth, and preprocessing steps. More explicit measurement-error models could improve the treatment of zeros and low-abundance observations.

A further limitation is that the pathway ranking should be interpreted as exploratory rather than causal. Cross-entropy analysis identifies pathways that are informative for time or phase prediction, but it does not prove that these pathways directly cause sensory spoilage. Experimental validation would be needed to confirm whether the identified modules are mechanistically involved in the production of spoilage-associated metabolites. Integrating metatranscriptomic trajectories with volatile organic compounds, metabolomics, microbial abundance data, and sensory scores would help connecting functional pathway activity more directly to the biochemical causes of off-odors and product rejection.

Future work should therefore focus on external validation, richer experimental designs, and multi-omics integration. A natural next step would be to train models across several storage temperatures and production lots, allowing the model to learn both shared spoilage trajectories and condition-specific differences in progression rates. Another direction would be to combine functional pathway trajectories with sensory and volatile compound data to identify which transcriptional changes are most closely linked to the compounds responsible for spoilage perception. Finally, the most robust pathway modules identified across temperatures could be evaluated as candidate biomarkers for targeted assays or predictive shelf-life tools.

In conclusion, this study demonstrates that longitudinal metatranscriptomic pathway profiles contain a strong and biologically interpretable signal of MAP broiler meat spoilage progression. The censoring-aware Gaussian process framework predicted storage time accurately within the 4°C experiment, produced predictions aligned with sensory spoilage phases, and revealed that selected aspects of the functional progression observed at 4°C were also detectable in the 6°C series within the investigated experimental system. These findings support the use of probabilistic time-series modeling to move shelf-life research beyond retrospective microbial enumeration toward a predictive, function-based understanding of spoilage dynamics.

## Supporting information

Supplemental analyses

cross-entropy scores 6 deg

cross-entropy scores 4 deg

## 5. SUPPLEMENTARY MATERIAL

Supplementary Material File 1: sup-file-cross-entropy-scores-per-pwy-per-tp-4-deg.xlsx Supplementary Material File 2: sup-file-cross-entropy-scores-per-pwy-per-tp-6-deg.xlsx Supplementary Material File 3: sup-sensitivity-analyses.pdf

## 6. DATA AVAILABILITY STATEMENT

The RNA-Seq data were deposited in NCBI SRA under Bioproject number PRJNA1464459 and are publicly available.

## 7. CODE AVAILABILITY

The Python code for implementing our method is publicly available and can be found on https://github.com/TrustworthyMLHelsinki/censoring-aware-GP.git

