## Supplemental analyses for "Censoring-aware Gaussian process modeling identifies time informative functional dynamics associated with spoilage status of modified atmosphere packaged broiler meat"

### Supplementary Material

#### S.1 Sensitivity analysis for $\tau$ and $\sigma_{\text{missing}}$

Table ?? shows the sensitivity analysis for different choices of $(\tau, \sigma_{\text{missing}})$, how they affect the LORO analysis shown in the main paper.

Table 1: Leave one replicate (LORO) out analysis for different values of $\tau$ and $\sigma_{\text{missing}}$. In each case the absolute mean error (MAE) is reported.

| Setting | $\tau$ | $\sigma_{\text{missing}}$ | MAE |
|---|---|---|---|
| Estimated from data $(\tau, \sigma_{\text{missing}})$ | 51.2 | 1.83 | 0.434 |
| $(0.5 \times \tau, \sigma_{\text{missing}})$ | 25.6 | 1.83 | 0.934 |
| $(1.5 \times \tau, \sigma_{\text{missing}})$ | 76.8 | 1.83 | 0.503 |
| $(\tau, 0.5 \times \sigma_{\text{missing}})$ | 51.2 | 0.915 | 0.434 |
| $(\tau, 1.5 \times \sigma_{\text{missing}})$ | 51.2 | 2.745 | 0.462 |

#### S.2 Sensitivity analysis for top $k = \{25, 50, 100\}$, highest ranking pathway modules according to cross-entropy scores

Table ?? shows the overlap significance among the top (k) pathway–taxon entries (according to cross-entropy scores), for the pathway–taxon resolution and the broader module resolution.

Table 2: Permutation-based sensitivity analysis of pathway–taxon and module overlap between the 4 °C and 6 °C datasets. The null mean and 95% interval were obtained from the permutation distribution.

| Phase | Level | Top $k$ | #pathw. 4 °C | Eligible 6 °C | Shared pathw. | Shared mod. | Obs. overlap | Null mean | Null 95% interval | Fold enrich. | $p_{\text{emp}}$ | $q_{\text{BH}}$ |
|---|---|---|---|---|---|---|---|---|---|---|---|---|
| Fresh prod. | Pathw-taxon | 25 | 362 | 143 | 142 | 134 | 7 | 1.71472 | [0, 4] | 4.082 | $7.40 \times 10^{-4}$ | $2.22 \times 10^{-3}$ |
| Fresh prod. | Module | 25 | 362 | 143 | 142 | 134 | 10 | 4.19632 | [1, 8] | 2.383 | $1.76 \times 10^{-3}$ | $4.53 \times 10^{-3}$ |
| Fresh prod. | Pathw-taxon | 50 | 362 | 143 | 142 | 134 | 9 | 6.85337 | [3, 11] | 1.313 | 0.2268 | 0.2721 |
| Fresh prod. | Module | 50 | 362 | 143 | 142 | 134 | 15 | 15.46723 | [10, 21] | 0.970 | 0.6353 | 0.6727 |
| Fresh prod. | Pathw-taxon | 100 | 362 | 143 | 142 | 134 | 32 | 27.42235 | [20, 35] | 1.167 | 0.1440 | 0.1994 |
| Fresh prod. | Module | 100 | 362 | 143 | 142 | 134 | 46 | 52.57610 | [46, 59] | 0.875 | 0.9777 | 0.9777 |
| Early sp. | Pathw-taxon | 25 | 402 | 143 | 143 | 135 | 3 | 1.55385 | [0, 4] | 1.931 | 0.1962 | 0.2523 |
| Early sp. | Module | 25 | 402 | 143 | 143 | 135 | 8 | 4.10873 | [1, 8] | 1.947 | 0.0270 | 0.0541 |
| Early sp. | Pathw-taxon | 50 | 402 | 143 | 143 | 135 | 11 | 6.22676 | [2, 11] | 1.767 | 0.0311 | 0.0559 |
| Early sp. | Module | 50 | 402 | 143 | 143 | 135 | 15 | 15.19925 | [10, 21] | 0.987 | 0.5970 | 0.6717 |
| Early sp. | Pathw-taxon | 100 | 402 | 143 | 143 | 135 | 40 | 24.86847 | [18, 32] | 1.608 | $7.00 \times 10^{-5}$ | $3.60 \times 10^{-4}$ |
| Early sp. | Module | 100 | 402 | 143 | 143 | 135 | 58 | 51.92488 | [45, 59] | 1.117 | 0.0579 | 0.0868 |
| Late sp. | Pathw-taxon | 25 | 403 | 143 | 143 | 135 | 7 | 1.54795 | [0, 4] | 4.522 | $3.90 \times 10^{-4}$ | $1.40 \times 10^{-3}$ |
| Late sp. | Module | 25 | 403 | 143 | 143 | 135 | 9 | 4.11959 | [1, 8] | 2.185 | $8.05 \times 10^{-3}$ | 0.0181 |
| Late sp. | Pathw-taxon | 50 | 403 | 143 | 143 | 135 | 16 | 6.20456 | [2, 11] | 2.579 | $5.00 \times 10^{-5}$ | $3.60 \times 10^{-4}$ |
| Late sp. | Module | 50 | 403 | 143 | 143 | 135 | 26 | 15.21348 | [10, 21] | 1.709 | $8.00 \times 10^{-5}$ | $3.60 \times 10^{-4}$ |
| Late sp. | Pathw-taxon | 100 | 403 | 143 | 143 | 135 | 59 | 24.82003 | [18, 32] | 2.377 | $1.00 \times 10^{-5}$ | $1.80 \times 10^{-4}$ |
| Late sp. | Module | 100 | 403 | 143 | 143 | 135 | 58 | 51.90637 | [45, 59] | 1.117 | 0.0563 | 0.0868 |

The empirical $p$-value ($p_{\text{emp}}$) quantifies the probability of obtaining an overlap at least as large as the observed overlap under the permutation null distribution. The $q_{\text{BH}}$ values are Benjamini–Hochberg-adjusted $p$-values calculated across all tests.
